# Development and Genome-level Microevolution of Oral Microbiome during Surface Colonization

**DOI:** 10.1101/2025.11.07.685765

**Authors:** Yuchen Zhang, Yuguang Yang, Yibing Liu, Emily Ming-Chieh Lu, David Moyes, Sadia Ambreen Niazi, Qin Zhou

## Abstract

The oral microbiome is essential to human health, yet its *de novo* ecological succession and microevolutionary dynamics remain poorly understood due to the absence of tractable *in vivo* models. Dental implants provide a unique opportunity to investigate these processes by establishing a defined time-zero for initial bacterial attachment. In this cohort study, we performed shotgun metagenomics on 95 subgingival plaque samples from 19 participants, including peri-implant sites at weeks 1-4 after crown placement and adjacent teeth as controls. Peri-implant and adjacent periodontal microbiomes exhibited distinct taxonomic and functional profiles. Even the same taxa had different functional potential across the two sites. These findings indicated that oral microbiome development is a niche-specific process with selective colonization rather than passive microbial translocation from adjacent sites. Longitudinal analysis identified three microbial community modules driving the succession of oral microbiome, including pioneer colonizers, constitutive species, and late commensals. Each module followed distinct temporal abundance patterns and played unique ecological roles throughout the succession process. Strain-level resolution revealed divergent microevolutionary trajectories: pioneer colonizers and late commensals exhibited higher cumulative mutation rates and greater strain heterogeneity overtime, with enrichment of nonsynonymous single nucleotide variants in genes related to virulence and metabolism, whereas constitutive species remained evolutionarily stable by contrast. Our study reveals that the development, succession, and microevolution of the oral microbiome is structured, niche-dependent, and modulated by inter-species facilitation and selective genomic adaptation. These findings advance the understanding of oral microbiome ecology and provide a conceptual foundation for manipulating microbial succession in health and disease.

## Introduction

The oral cavity is a highly diverse, dynamic ecosystem comprised of multiple spatially distinct habitats [1]. These include the hard, non-shedding surfaces of natural teeth and prosthetic materials (e.g. dentures, crowns, implants), the soft epithelial surfaces of the oral mucosa (e.g. buccal and gingival mucosa), and additional niches such as tongue dorsum and gingival sulcus [2]. Each habitat exhibits unique histological and spatial features resulting in distinct physicochemical microenvironments (pH, oxygen tension, and nutrition availability) thereby shaping site specific microbial communities [3, 4].

Oral microbial communities are complex systems regulated by dense microbe-microbe networks, constant host-microbe interactions, and recurrent external perturbations including dietary intake, salivary flow, and oral hygiene procedures. These factors repeatedly shift the local conditions, impacting the selective pressures experienced by individual microbes [5, 6]. Together, these elements contributed to the uniqueness and complexity of the oral microbiome. Within this complexity, the microbial communities continually adjust themselves in response to environmental perturbations, host immune regulation, and changes in selective pressure [7]. These responses manifest as both compositional changes and functional alterations [3]. Genome-resolved longitudinal studies show rapid, selection-driven microevolution *in vivo*, altering gene content and functional potential via selective sweeps, single nucleotide variances (SNVs), and horizontal gene transfer, even under homeostatic conditions [8–10].

Currently, most literature has focused on established communities in the oral cavity, investigating how the oral microbiome maintains homeostasis and how dysbiosis emerges during diseases [11, 12]. By contrast, fewer studies have evaluated *de novo* community assembly and ecological dynamics in oral microenvironments, including different ecological roles of oral taxa and their microevolution trajectories. Conventionally, the formation of new oral microbial communities has been regarded primarily as a passive consequence of bacterial migration from adjacent habitats. However, emerging evidence suggests that *de novo* community assembly should be far more complex, involving structured ecological succession, selective pressures imposed by local environment, and potential microevolution that shapes community composition and function [13].

Due to practical and ethical constraints, investigations of *de novo* community assembly on natural surfaces in the oral cavity are difficult to conduct. In this scenario, dental implants offer a feasible solution. Their supra-structures, including abutments and crowns, are fully sterilized before being placed into the mouth, providing a defined time point for initial exposure to the microenvironment [14]. Biofilms then start to establish within hours to days, enabling longitudinal investigation on this process [15]. The surfaces of the implant components function as standardized, hard, non-shedding surfaces with well-defined topography. Together, these features make implants a tractable *in vivo* model for studying *de novo* community assembly and microevolution in the complex oral microenvironment. Here, we conducted a longitudinal cohort study to track peri-implant microbiome development from the day of final abutment and crown placement up to week four, using shotgun metagenomic sequencing. The adjacent periodontal microbiome served as the control. We aimed to determine whether community assembly of oral microbiome is consequential to the bacterial translocation from adjacent niches, and to characterize the development of the oral microbiome in complex microenvironments together with genome-level microevolution within its community.

## Methods

### Ethics approval

This study was approved by the Ethics Committee of Xi’an Jiaotong University (No. 2023-XJKQIEC-025-002). The performance of this study adhered to the Strengthening the Reporting of Observational Studies in Epidemiology (STROBE) guidelines. Written consent was obtained from each participant.

### Participant recruitment and sample collection

This study recruited 19 patients with a single missing tooth that required implant restoration. The patients were free of other oral conditions. All implants used were commercial titanium bone-level implants. Detailed inclusion and exclusion criteria are described in Table S1. The surgical stages were performed by an experienced clinician following a standard protocol of implant placement with submucosal healing. After 3 to 6 months of healing, cone beam computed tomography (CBCT) scans were taken to confirm radiographic evidence of implant osseointegration. At the second-stage surgery, healing abutments were installed to shape the soft and hard tissue around the implant neck, followed by the placement of restoring abutments and the final crown.

Subgingival plaque samples from the implant were collected at 1 week, 2 weeks, 3 weeks, and 4 weeks after final crown placement. As a control group, subgingival plaque samples from the adjacent teeth were also collected at 1 week (Fig. 1). Prior to sample collection, participants were first asked to rinse their mouth with distilled water for 10 s. The selected sites were isolated using sterile cotton rolls to avoid contamination from the saliva. The supragingival plaque was gently removed with cotton pellets. The subgingival plaque samples were collected by inserting sterilized endodontic paper points (Dentsply Sirona, Bensheim, Germany) into the gingival sulcus for 30 seconds. This step was repeated a total of 6 times to ensure sufficient absorption of sulcus fluid and attachment of microbes. The collected paper points were placed in 1.5 mL microcentrifuge tubes (Eppendorf, Hamburg, Germany) containing phosphate buffered saline and frozen at -80 °C for further use. All samples underwent DNA extraction within 4 weeks of collection.

**Fig. 1.**
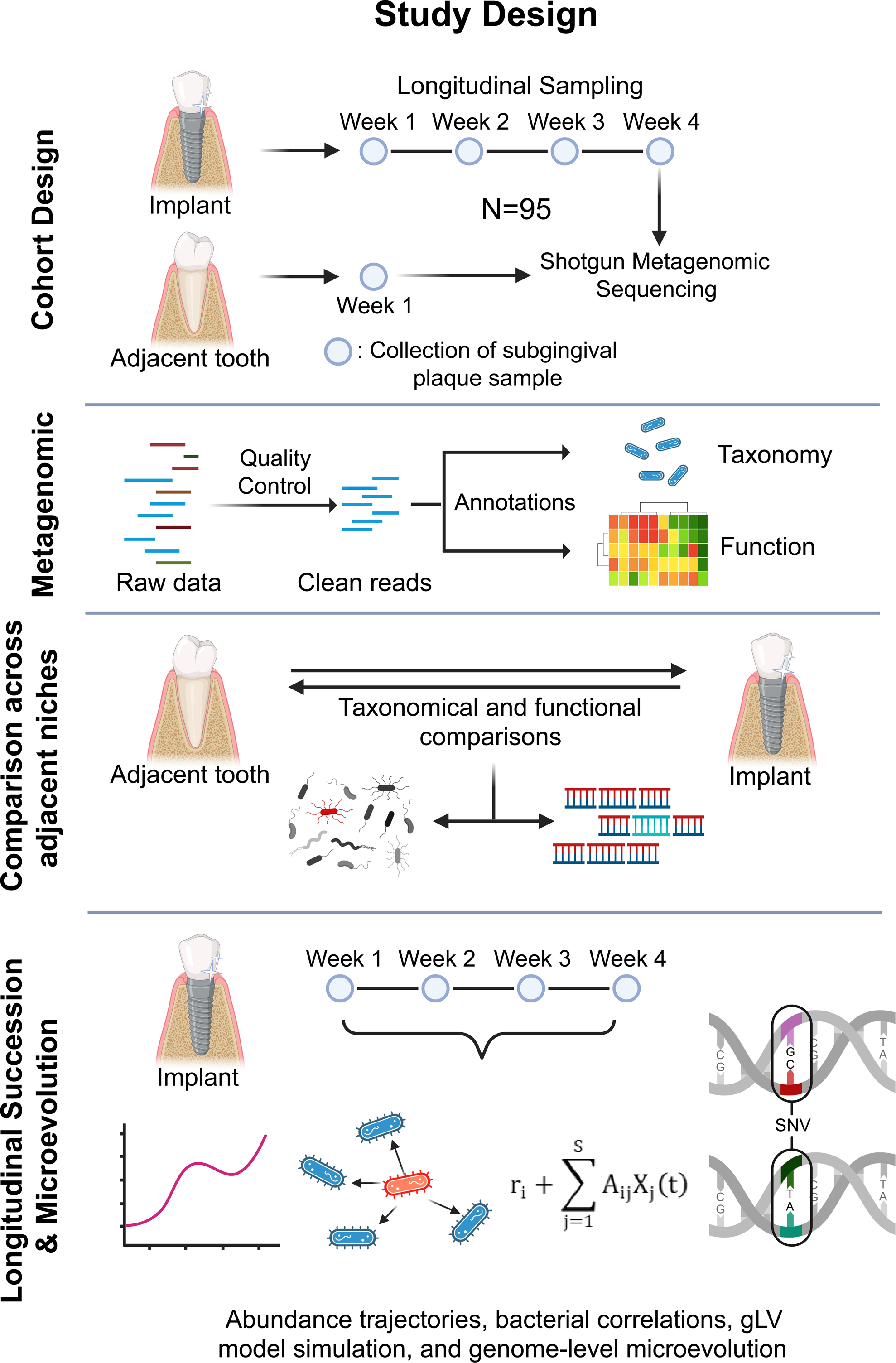
A flowchart of study design.

### DNA extraction and metagenomic sequencing

Bacterial DNA was extracted from subgingival plaque samples using E.Z.N.A. Soil DNA kits (Omega Bio-tek, USA) following the instructions of the product manual. The extracted DNA was then quantified using Qubit dsDNA HS assay kits (Thermo Fisher Scientific, USA). Samples with sufficient DNA yield were then subjected to metagenomic library preparation and were pair-end sequenced using an Illumina NovaSeq 6000 PE150 (Illumina, CA, USA).

### Sequence quality control

The raw sequences were quality-controlled and decontaminated using *KneadData* [16]. Briefly, the adapters and low-quality bases were trimmed using *Trimmomatic* with a sliding window cutoff of 4:20 and a minimum read length of 50 bp. The trimmed reads were then aligned against the human genome reference (hg39) using *Bowtie2* under very-sensitive mode to remove host-derived contaminant sequences. Only reads that passed quality control on both directions were retained for downstream analyses. The quality of the cleaned reads was then checked with *FastQC* to ensure sufficient trimming and decontamination.

### Taxonomic and functional annotations of the microbiome reads

*MetaPhlAn4* was used to align the cleaned reads against *ChocoPhlAn4* database to generate taxonomic annotations [17]. Per-sample relative abundances at the species level were extracted for downstream statistical analyses. *HUMAnN3* was employed to detect the functional capacity of the microbiomes with *ChocoPhlAn4* database and UniRef90 protein reference [18]. The identified pathways were regrouped into metabolic pathways based on the MetaCyc database and normalized to copies per million (CPM) unit for downstream analyses.

### Alpha and beta diversity analysis

The alpha and beta diversity of each sample was calculated using *vegan* package in R (version 4.4.1). Alpha diversity was evaluated using Shannon and Simpson indices. Beta diversity was evaluated based on Bray-Curtis distance and was visualized using Principal Coordinate Analysis (PCoA).

### Identification of differentiating species and pathways

To identify the bacterial species and gene pathways that had significantly higher abundance in the peri-implant microbiome compared to periodontal microbiome (namely differentiating species and pathways), *MaAsLin3* was used to compare the prevalence (logistic models) and abundance (linear models) of each annotated species or pathway between peri-implant and periodontal microbiomes, adjusting for read depth, gender, age, and implant brand as fixed effects [19]. Bacterial relative abundance was normalized using total-sum scaling (TSS) and log-transformation prior to comparisons, while pathway abundance was compared in CPM units.

### Species abundance heatmap and hierarchical clustering

The Z-scored relative abundance of the top 50 abundant species across the samples and time points were visualized as a heatmap using *ComplexHeatmap* R package. These species were then hierarchically clustered into three community modules based on Euclidean distance and complete linkage, as implemented in *ComplexHeatmap*. For each module-time stratum, the top 5 abundant pathways were also visualized.

### Computation of bacterial co-occurrence network

To compute the bacterial co-occurrence networks, the correlations among bacterial species in peri-implant microbiome were tested using *psych* package in R based on Spearman correlation coefficients. The p-values were adjusted by controlling False Discovery Rate (FDR). Correlations with a |rho| > 0.4 and p-FDR < 0.05 were extracted to construct the co-occurrence network for each time point.

### Simulation analysis of attachment probability

To assess whether the attachment of late commensals was facilitated by the presence of constitutive species, we analysed their attachment probabilities through generalized Lotka–Volterra (gLV) model using *MATLAB* (r2024b). We assume a resident community composed of *S* species at a feasible equilibrium 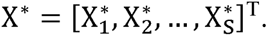 Then based on gLV model, the dynamics of the resident species are described by

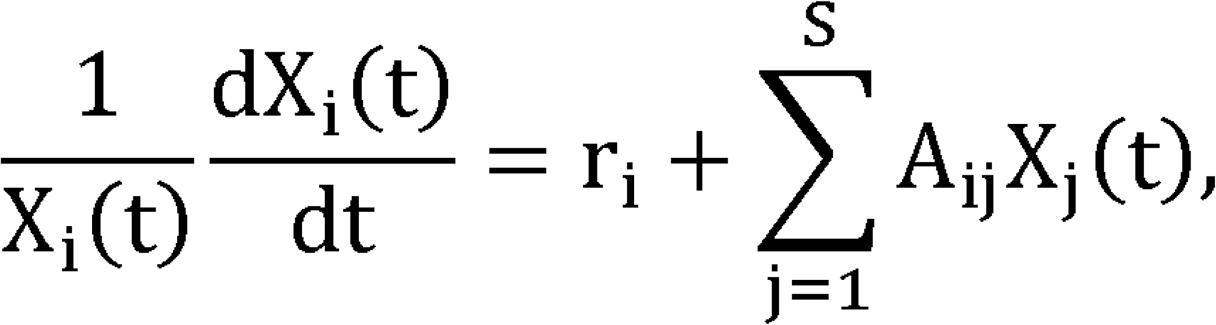

where *X_i_*(*t*) denotes the abundance of species *i* at time *t*, and *A_ij_* captures the effect of species *j* on species *i*. A feasible equilibrium satisfies

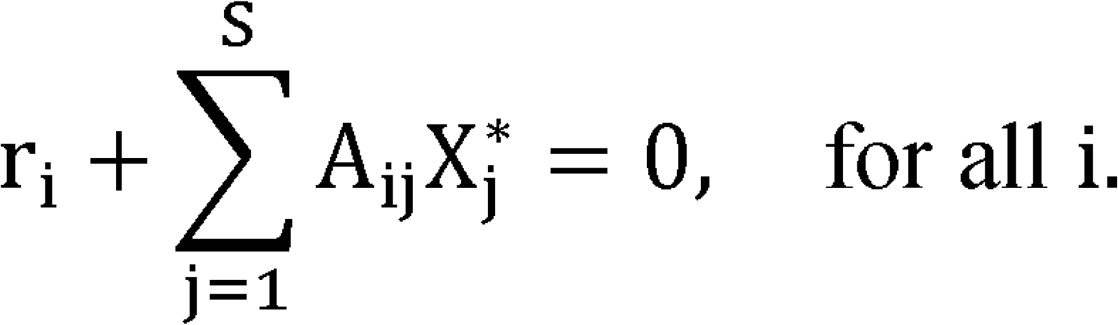

When a new species (indexed as *S* + 1) is introduced at a low abundance, its per capita growth rate is given by

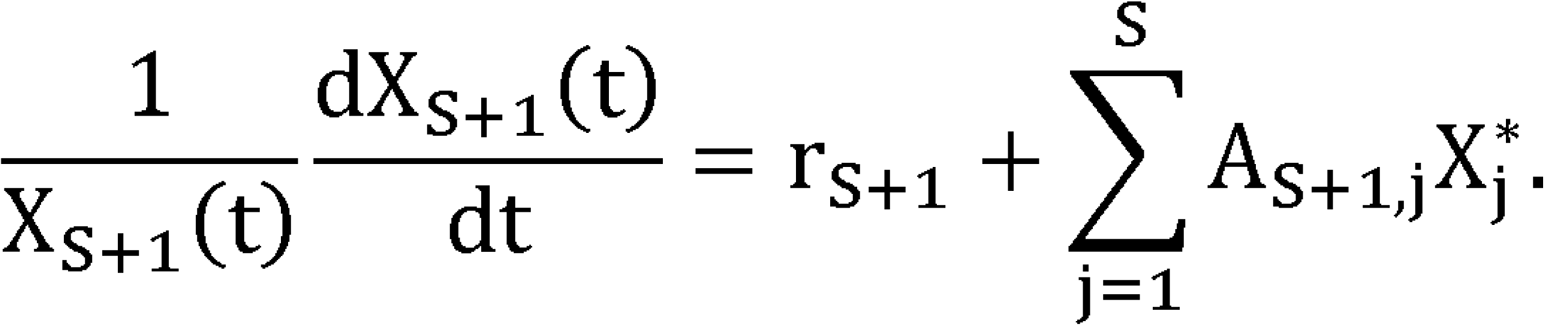

If the right-hand side of this equation is positive, the species can grow when rare and is therefore considered able to attach. This condition is considered as the attachment criterion, similar to invasion criterion reported in other studies [20, 21].

Within this framework, we constructed two dummy communities, one with pioneer colonizers only (P) and one with both pioneer colonizers and constitutive species (P + C). The interaction types among the bacterial species were inferred from the co-occurrence networks computed above, and the strengths of the interactions were sampled from a random distribution. We performed 1000 simulations for each late commensal species and denoted the proportion of simulations that met the attachment criterion (*N*) as attachment probability (*N*/1000).

### Detection of strain-level mutations and strain heterogeneity

*StrainPhlAn* was employed to detect the strain-level mutations of the top 50 most abundant species [22]. The consensus marker sequences of each species were extracted according to their species-level genome bins (SGBs) and were reconstructed. Sample sequences were aligned against the reconstructed marker sequences, allowing identification of temporal nucleotide variations within the same species. Samples were filtered with a minimum of 20 marker genes per sample and a 25% marker presence threshold prior to downstream analysis. Only markers present in at least 50% of primary samples were retained. If the marker genes were detected at more than two time points including week 1 in a given subject, the samples of that subject will be included for longitudinal comparisons for cumulative mutation rates.

*InStrain* was further used to assess the strain-level single nucleotide variants (SNV) and strain heterogeneity across the time points in the selected species [23]. Briefly, the cleaned reads were mapped against a customized database comprising reference genomes (NCBI RefSeq) of the selected species. For each species, inStrain detected its SNVs (compared to reference genome) from week 1 to 4 and determined if these SNVs were synonymous, non-synonymous, or intergenic with Prodigal prediction [24]. Non-synonymous to synonymous SNV ratio (N/S ratio) was calculated for each species and were compared longitudinally overtime. Temporal strain heterogeneity was evaluated based on population average nucleotide identity (popANI) and shared genome coverage (SGC). A threshold of 99.9% popANI was used as an indicative level of potential strain shifts.

For the non-synonymous SNVs detected above, we also examined the corresponding KEGG Orthology (KO) terms of the genes in which these SNVs were located. The KO terms of the references genome and sample sequences were annotated using KoFamScan. We then aggregated the number of non-synonymous SNVs per KO across samples and species. For each species at each time point, the top 5 KO terms that were most frequently affected by non-synonymous SNVs were extracted. Enrichment analyses based on Fisher’s exact tests were subsequently performed to check if these KOs were disproportionately impacted by non-synonymous SNVs in specific species. Through this, we were able to show potential directions of microbial functional shaping driven by microevolution.

### Statistical analysis

Unless otherwise specified, the statistical comparisons across the groups, including weeks 1 to 4 of peri-implant microbiome and the periodontal microbiome from adjacent teeth (AT), were conducted using Friedman tests with Dunn’s post-hoc multiple comparisons in GraphPad Prism (version 10.4.2).

## Results

### Demographics of the cohort

A total of 95 plaque samples were collected and analysed from 19 participants (Fig. 1), comprising peri-implant plaque at weeks 1 to 4 (n = 19 per time point), and adjacent periodontal plaque at week 1 (n = 19). The demographics of the participants, including age, gender, and implant brands were summarized in Table S2 and Data S1. The mean age of the cohort was 37.32 ± 11.22 years old. There were 14 (73.68%) female participants and 5 (26.32%) male participants. The included implants were from 5 manufacturers including Bego (42.11%), Straumann (31.58%), Osstem (5.26%), Hiossen (15.79%), and Zimmer (5.26%).

### Distinct bacterial composition at adjacent oral microenvironments

Following the metagenomic workflow, a total of 617 bacterial species were identified in these 95 samples. Among them, 299 species were shared by both peri-implant and periodontal ecological niches, whilst 41 and 51 species were exclusive to peri-implant and the adjacent periodontal microbiomes, respectively (Fig. 2A). Notably, within the peri-implant niche, weeks 1 to 4 each contained week-specific species in addition to a shared core, indicating ongoing community succession during biofilm assembly (Fig. 2A). Compositional differences between peri-implant and periodontal microbiomes were further confirmed by diversity analyses. Overall, peri-implant microbiome exhibited poorer microbial richness and evenness when compared to periodontal microbiome (Fig. 2B), with significantly lower Shannon index at weeks 1 and 2 (p < 0.05) and significantly lower inverse Simpson index at weeks 1 and 3 (p < 0.05). Principal coordinate analysis (PCoA) was used to visualize the beta diversity of the microbiomes based on Bray-Curtis distance (Fig. 2C). The microbial compositions of peri-implant microbiome were significantly different from adjacent periodontal microbiome (p < 0.0001, Kruskal-Wallis test on principle coordinate 1).

**Fig. 2.**
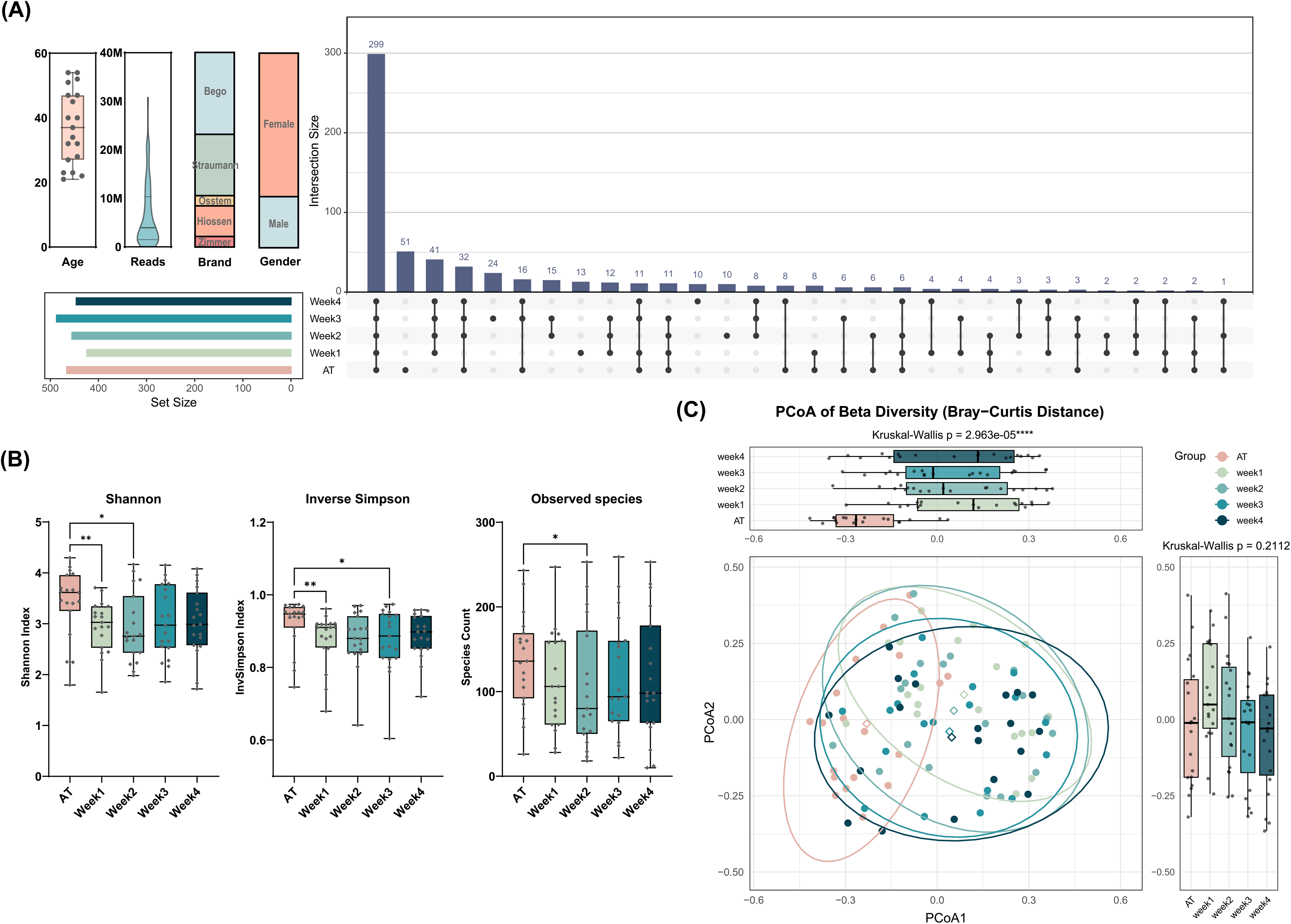
Distinct bacterial composition in peri-implant and periodontal microbiomes. **(A)** Demographic features of the cohort and an UpSet plot showing the number of bacterial species present in each group. **(B)** Comparison of alpha diversity based on Shannon index, inverse Simpson index, and the number of observed species. **(C)** Principal Coordinate Analysis (PCoA) of beta diversity based on Bray-Curtis distance. The ellipses represented the 90% confidence levels while the diamonds represented their centroids. Projections on principle coordinate 1 and 2 were compared amongst groups using K-W tests. * p < 0.05, ** p < 0.01, **** p < 0.0001, AT: adjacent teeth.

### Differential taxa and functions at adjacent oral microenvironments

To identify the bacterial species that were significantly different in either abundance or prevalence between peri-implant and periodontal microbiomes, *MaAsLin3* was utilized based on linear and logistic regression models respectively, while adjusting for read depth, gender, age, and implant brands (Fig. 3A). The results showed that *Streptococcus anginosus*, *Prevotella melaninogenica*, *Veillonella rogosae*, *Slackia exigua*, *Neisseria sicca*, *Rothia mucilaginosa*, and *Lancefieldella rimae* were the most significantly enriched species in peri-implant microbiome compared to the adjacent periodontal microbiome (Fig. S1). The abundance and prevalence of these peri-implant-enriched species were not significantly associated with confounding factors, with the exception of *N. sicca*, which was positively associated with male gender and Bego brand implants (Fig. 3A and Fig. S2). Among these 7 species, 6 of them demonstrated significantly higher relative abundance (p < 0.05, Friedman test with Dunn’s post-hoc) at one or more time points in peri-implant microbiome compared to the periodontal control (Fig. 3B). These species made up around 15-20% of the peri-implant community but less than 3% of the periodontal community (Fig. 3C), underlining niche-specific assembly of the oral microbiome.

**Fig. 3.**
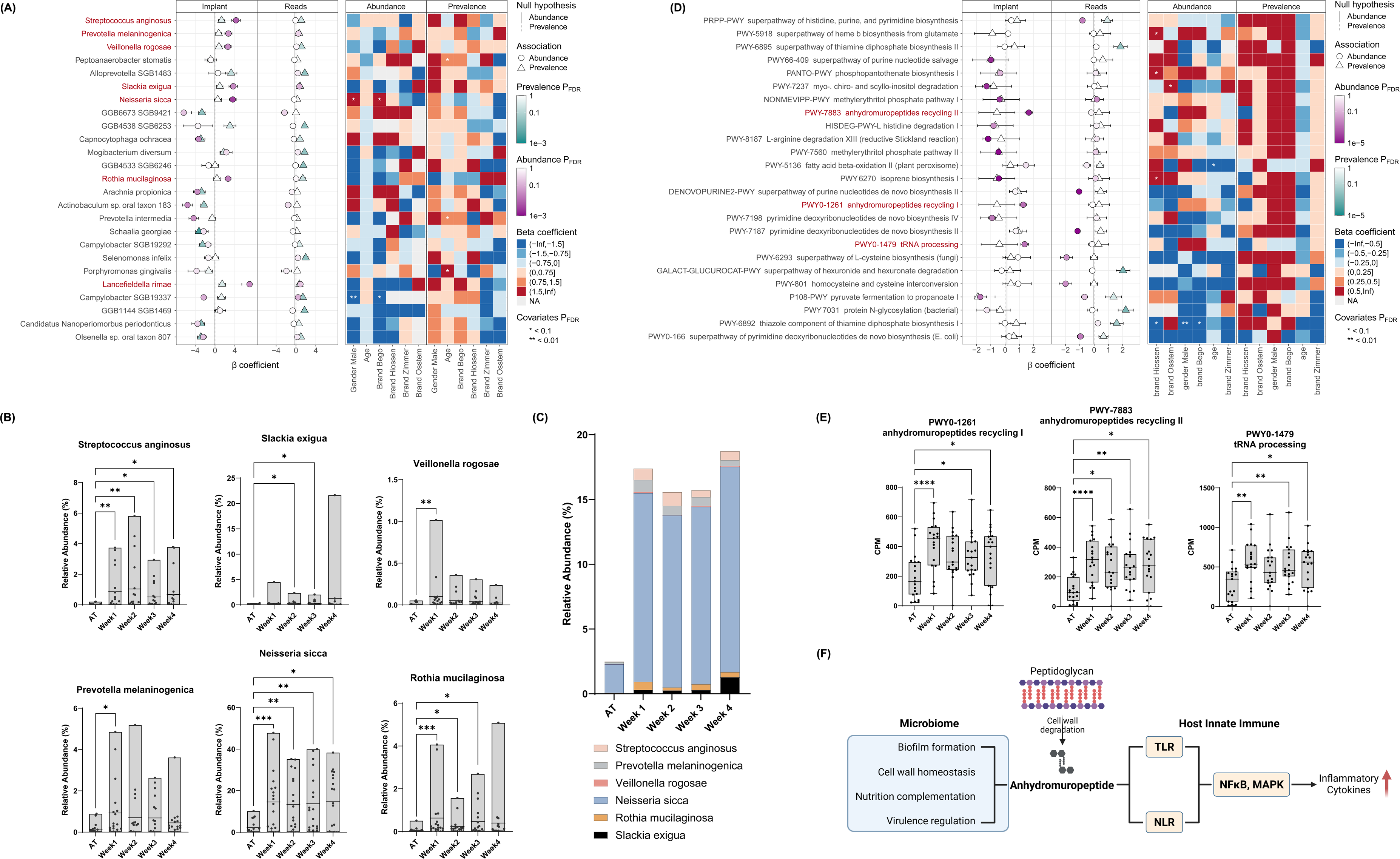
Peri-implant-enriched species and functional pathways. **(A)** Linear and logistic regression models of bacterial abundance and prevalence in peri-implant microbiome compared to adjacent periodontal microbiome, adjusting for read depth, gender, age, and implant brands. Species highlighted in red were significantly more abundant in peri-implant microbiome (adjusted p-value < 0.05), namely peri-implant-enriched species. **(B)** Relative abundance of peri-implant-enriched species. **(C)** The proportion (summed relative abundance) of peri-implant-enriched species in peri-implant and adjacent periodontal microbiomes. **(D)** Linear and logistic regression models of pathway abundance and prevalence in peri-implant microbiome compared to adjacent periodontal microbiome, adjusting for read depth, gender, age, and implant brands. Pathways highlighted in red were significantly more enriched in peri-implant microbiome (adjusted p-value < 0.05). **(E)** Gene abundance (copies per million, CPM) of the peri-implant-enriched pathways in peri-implant and adjacent periodontal microbiomes. **(F)** The roles of anhydromuropeptides in both microbiome and host innate immune. * p < 0.05, ** p < 0.01, **** p < 0.0001.

Following a similar protocol, the functional pathways that were significantly enriched in peri-implant microbiome compared to adjacent teeth were also visualized (Fig. 3D & 3E). Pathways related to anhydromuropeptides recycling (PWY-7883 and PWY0-1261) and tRNA processing (PWY0-1479) had significantly higher abundance in peri-implant microbiome after adjustment for read depth, gender, age, and implant brands (Fig. S3). Anhydromuropeptide has multiple roles in the microbial community in biofilm formation, nutrition complementation, and virulence regulation. It is also involved in inflammation by triggering innate immune responses through toll-like receptors (TLRs) and NOD-like receptors (NLRs) signaling [25–27] (Fig. 3F).

The distinct bacterial compositions and functional capacities between peri-implant and its adjacent microbiomes indicated that the *de novo* assembly of oral microbiome is a niche-specific assembling process rather than passive translocation from neighboring sites.

### Niche-specific functional potential at species level

Although species differ at the different ecological niches, it is possible that their functional potentials remain unchanged at different sites. To address this, we next compared the functional potentials of peri-implant-enriched species between the two ecological niches. These species exhibited specific pathway enrichment or depletion in peri-implant microbiome when compared to the same species in adjacent periodontal microbiome (Fig. 4). Pathways that were differentially abundant between peri-implant and periodontal microbiomes were extracted and categorized into 5 main categories based on their functions: cell wall and extracellular matrix, energy metabolism, nucleotide metabolism, amino acid metabolism, and all others. In the peri-implant microbiome, *P. melaninogenica* exhibited pathway enrichment in the cell wall and extracellular matrix as well as amino acid metabolism categories, whilst *V. rogosae* showed enrichment in pathways related to nucleotide metabolism. *R. mucilaginosa*, *S. anginosus*, and *N. sicca* also demonstrated significant differences in gene abundances across several pathways between peri-implant and periodontal microbiomes (Data S2). These findings suggested niche-specific functional rewiring: even the same species can possess distinct functional potential across adjacent ecological niches.

**Fig. 4.**
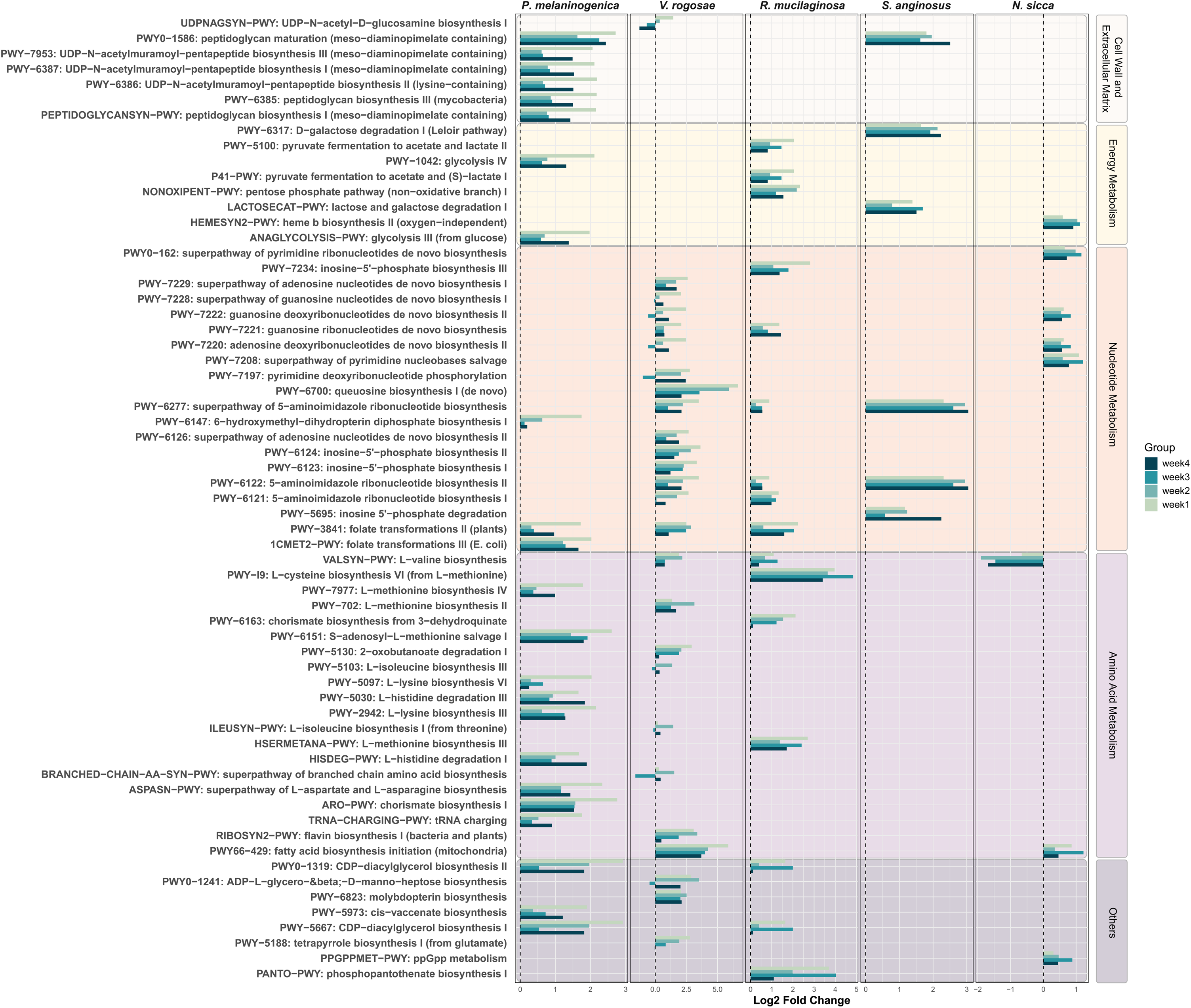
Functional differences of the same species across adjacent ecological niches. Functional pathways in peri-implant-enriched species that had significantly different abundance in the peri-implant microbiome (bars) compared to adjacent periodontal microbiome (dashed lines).

### Oral microbiome succession is driven by distinct microbial community modules

Having determined the species and functional differences between the two adjacent niches, we next investigated the succession dynamics of the peri-implant microbiome through longitudinal data series analysis. Firstly, the top 50 most abundant species were hierarchically categorized into three community modules according to the Euclidean distance of their Z-scored relative abundance across the time points (Fig. 5A and Data S3). Based on the trends in their relative abundance from week 1 to 4, these community modules were named as pioneer colonizers (M1), constitutive species (M2), and late commensals (M3). The pioneer colonizers were more abundant at weeks 1 and 2 than weeks 3 and 4 (Fig. 5A & 5B). In contrast, the late commensals exhibited the inverse temporal pattern, being less abundant at weeks 1 and 2 but more abundant at weeks 3 and 4 (Fig. 5A & 5B). The abundance of constitutive species remained stable showing no significant changes across all time points (Fig. 5A & 5B). The changes in module abundance were confirmed using Friedman tests with Dunn’s post-hoc comparisons (Fig. 5B).

**Fig. 5.**
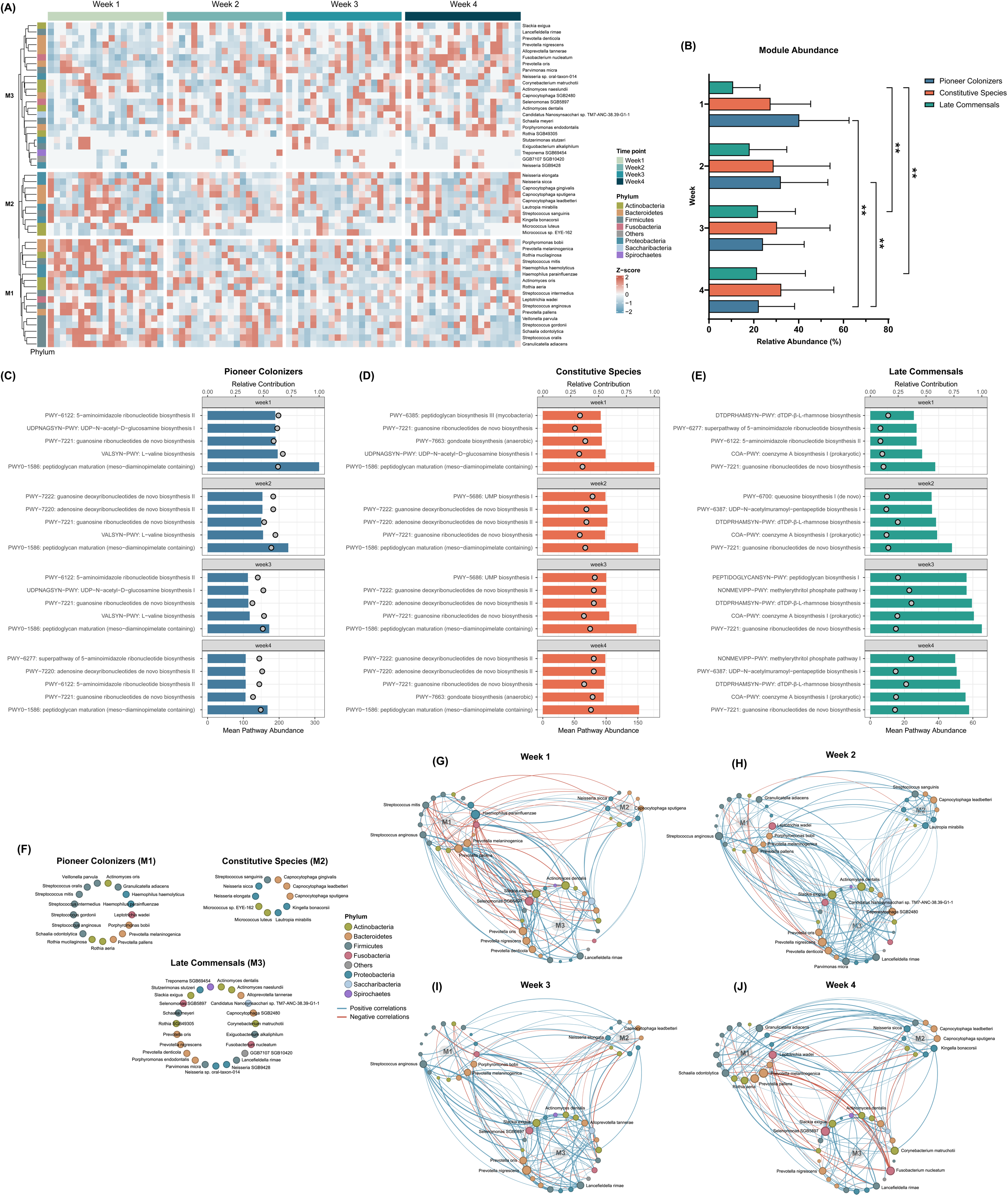
Oral microbiome succession driven by three microbial community modules. (A) Per-sample abundance (Z-score normalized) of the top 50 abundant bacterial species. Each column represents a sample while each row represented a bacterial species. (B) Module relative abundance across time points (Friedman test with Dunn’s post-hoc). (C-E) Top 5 abundant pathways in each module from week 1 to 4. Bars represent mean pathway abundance whilst circles represent relative contribution. (F) Members of each module and their corresponding phyla. (G-J) Co-occurrence networks of top 50 abundant species based on Spearman correlation coefficients. Each circle represents a bacterial species. Lager circles represent higher relative abundance. Blue and red lines represent positive and negative correlations, respectively.

To investigate if species from different modules were also distinct in their functional potentials, the top 5 enriched pathways in each module were extracted for each time point (Fig. 5C–5E and Data S4). In pioneer colonizers, these pathways were associated with bacterial cell wall and nucleotides metabolism (Fig. 5C), suggesting their importance in biofilm formation and community growth. The constitutive species were also highly abundant in pathways related to cell wall and nucleotides, but in addition were also abundant in uridine monophosphate (UMP) and gondoate biosynthesis, which were potentially linked with nitrogen and carbon fluxes in the microbial community [28]. In contrast, the late commensals were characterised by greater abundance in pathways involved in dTDP-β-L-rhamnose and coenzyme A biosynthesis.

Based on the relative abundance and the top-abundant gene pathways of each module, we hypothesized that the pioneer colonizers were highly competitive during the initial stage of peri-implant biofilm formation and can adapt to a microenvironment with limited resources, which made them more dominant during week 1 and 2. The constitutive species, although being less competitive than pioneer colonizers, were also able to survive the initial succession stage while having the potential to modulate the microenvironment to facilitate the attachment for late commensals. In contrast, the late commensals were less competitive during week 1 and 2, but their relative abundance gradually increased with peri-implant microbiome succession.

To validate our hypothesis through microbial interactions, we next computed the bacterial co-occurrence networks for each time point based on Spearman coefficients to infer the ecological role of each community module (Fig. 5G–5J and Data S5). An obvious characteristic of the week 1 network was that there were numerous negative correlations between pioneer colonizers (M1) and the other modules (Fig. 5G), suggesting that their presence negatively affected the growth of other species, potentially by competing for the limited ecological niches and resources. At week 2 and 3 (Fig. 5H & 5I), the number of negative correlations decreased tremendously. There were more correlations both within and between constitutive species (M2) and late commensals (M3). These findings indicated that the relative abundances of these bacteria were either rising or declining in a coordinated manner, suggesting that the community was shifting from a strong competition towards a co-adaptation to the microenvironment. The week 4 network was characterized by the reemergence of negative correlations. In contrast to the week 1 network, most of the negative correlations by week 4 involved late commensals rather than pioneer colonizers (Fig. 5J). This might suggest that the late commensals had gained advantages over other modules, which explained their increased abundance at week 4. In addition, the number of correlations were more evenly distributed both across and within the three modules at week 4, indicating that the community was evolving towards a more balanced and mature status.

### Attachment of late commensals is facilitated by constitutive species

Based on the correlations calculated above, we next constructed two dummy peri-implant microbial communities for each time point – one with only pioneer colonizers present (community P) and the other with both pioneer colonizers and constitutive species present (community P + C) – to simulate the attachment probability of late commensals in peri-implant microbiome using generalized Lotka-Volterra (gLV) models (Fig. 6A & 6B). In many cases, late commensals demonstrated higher attachment probability in community P + C compared to community P (Fig. 6C–6F), suggesting a facilitative role of constitutive species towards the attachment of late commensals. To quantitatively evaluate this effect, we further calculated the odds ratio of attachment probability between the P and P + C communities for each late commensal from week 1 to 4 (Fig. 6G). Notably, some species, including *Capnocytophaga SGB2480* (weeks 1-4), *Corynebacterium matruchotii* (week 2), *Neisseria sp. oral taxon 014* (week 2), and *Schaalia meyeri* (week 4), displayed significantly increased attachment probabilities (log_10_ OR > 0.5, p < 0.05) with the presence of constitutive species (Data S6).

**Fig. 6.**
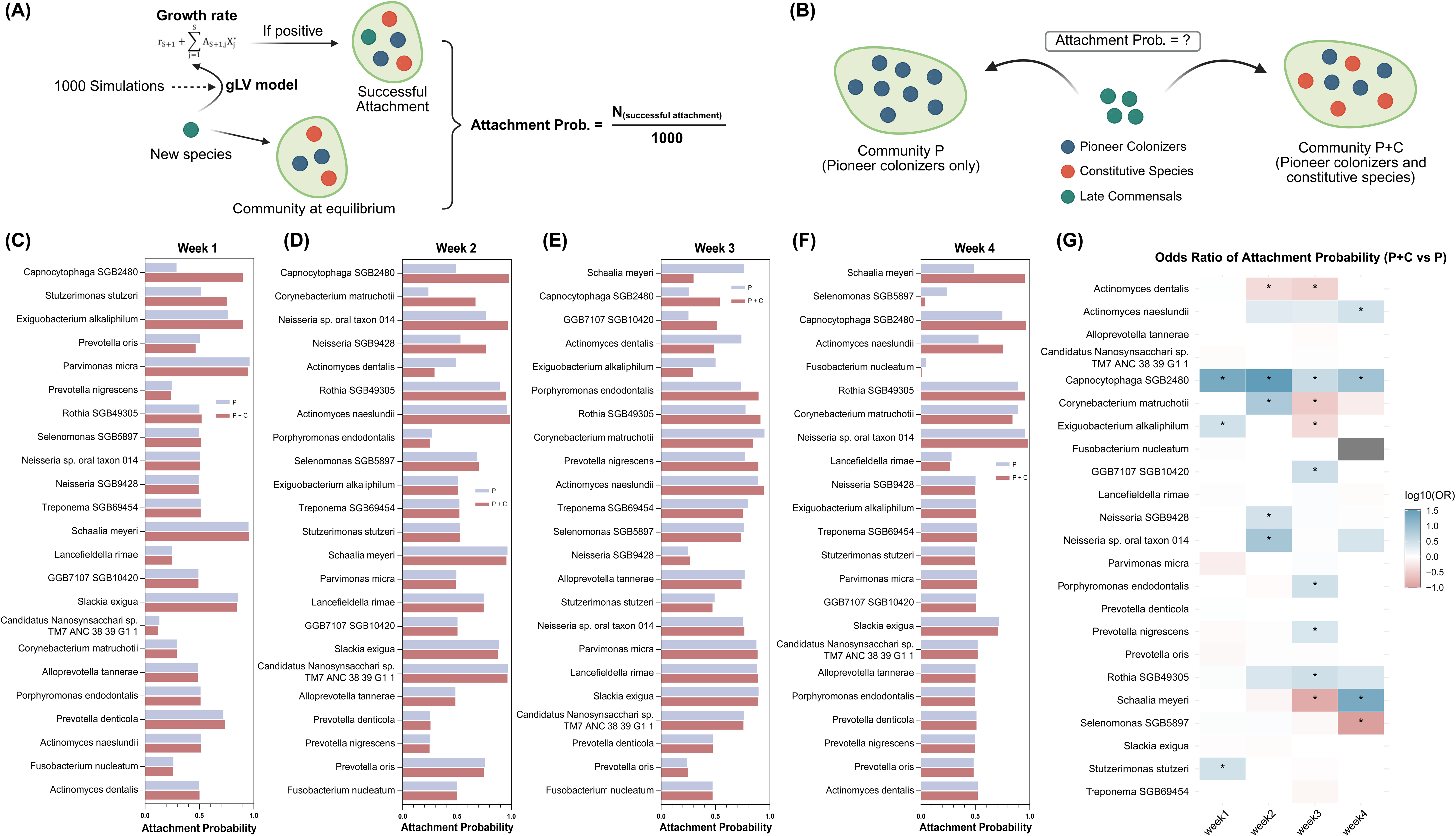
Attachment of late commensals is facilitated by constitutive species. **(A)** Generalized Lotka–Volterra (gLV) model used to predict attachment probability of the late commensals. Attachment of a newly introduced species is considered successful only when growth rate is positive. The attachment probability of this species is defined as the proportion of successful attachments out of 1000 simulations. **(B)** Attachment simulation on two dummy communities, one comprising only pioneer colonizers (community P) whilst the other consisting of both pioneer colonizers and constitutive species (community P + C). **(C-F)** Attachment probabilities of late commensals across weeks 1 to 4. Blue bars represent pioneer-only community (P) while red bars represent community with both pioneer colonizers and constitutive species (P + C). **(G)** Log_10_-transformed odds ratio of attachment probability between communities P + C and P. Positive values indicate higher attachment probability of late commensals in the presence of constitutive species, whereas negative values suggest potential inhibition. * p < 0.05.

### Strain-level heterogeneity and temporal microevolution in each community module

In order to see if different community modules also exhibited different extents of strain heterogeneity and microevolution, we compared longitudinally the consensus marker sequences of each species from week 1 to 4 using *StrainPhlAn*, and calculated their cumulative mutation rates at weeks 2 to 4 against week 1 (Fig. 7A). Overall, constitutive species exhibited lower cumulative mutations rates in their marker genes when compared to pioneer colonizers and late commensals (Fig. 7a and Fig. S4-S9). The differences at week 3 between constitutive species and both pioneer colonizers and late commensals, and at week 4 between constitutive species and late commensals were statistically significant (p < 0.05).

**Fig. 7.**
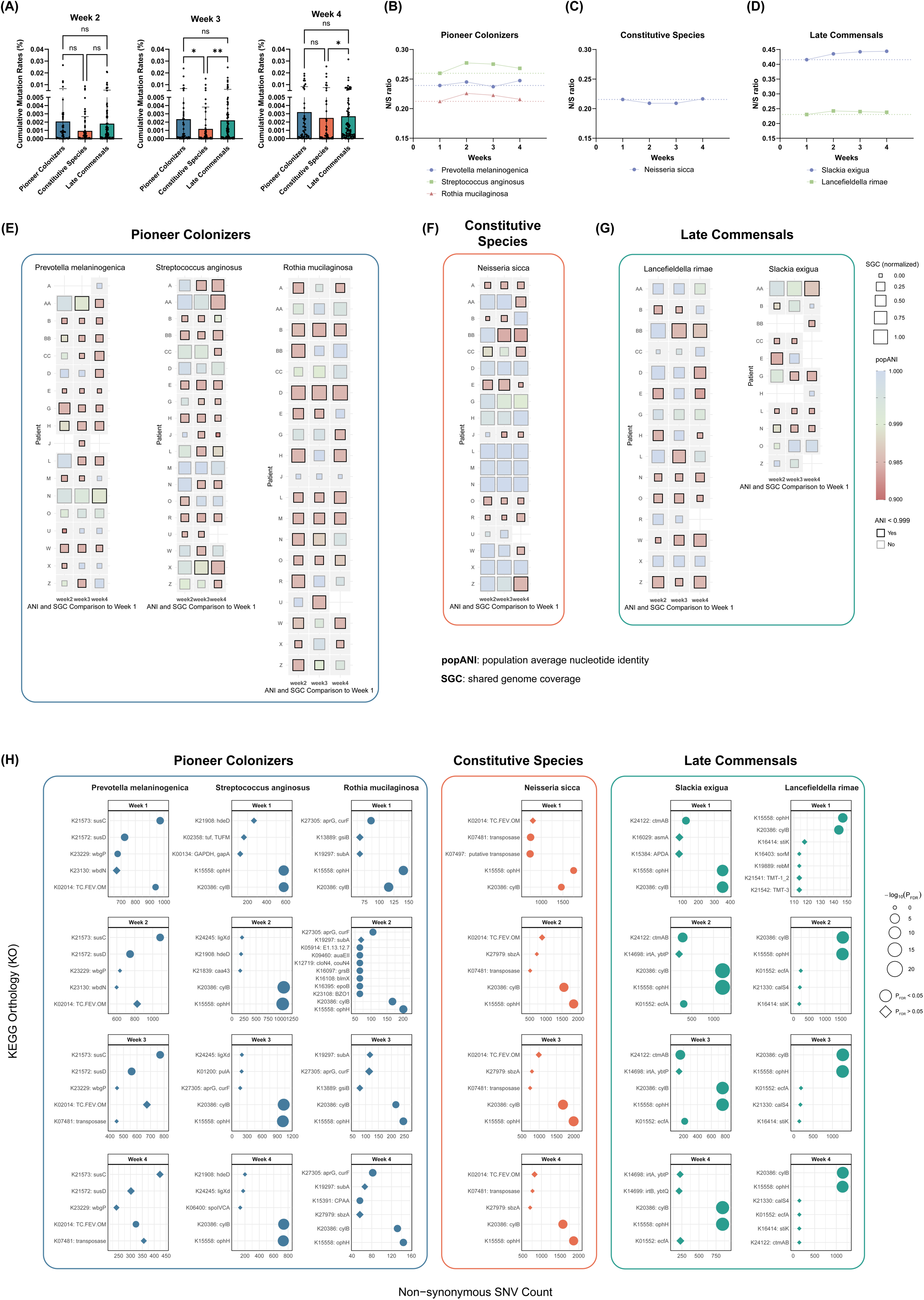
Assessment of temporal microevolution in each community module. (A) Cumulative mutation rates in consensus marker sequences at weeks 2 to 4 compared to week 1. * p < 0.05, ** p < 0.01. (B-D) Temporal alterations in the N/S ratios of the selected representative species from each community module. The dashed lines represent the week 1 N/S ratio level for each species. (E-G) Temporal strain heterogeneity in selected species assessed by popANI and SGC. Colour gradients reflect popANI values while square sizes reflect normalized SGC. Squares highlighted in black boarders denote popANI < 0.999, suggesting potential strain shifts. (h) Top KO terms with the highest counts of non-synonymous SNVs for each representative species and time point. Circles indicate statistically significant enrichment of non-synonymous SNVs (PFDR < 0.05), while diamonds indicate non-significant enrichment.

To assess the evolutionary pressure of different modules, we further looked into representative species of each community module (the peri-implant-enriched species identified above. *P. melaninogenica*, *S. anginosus*, and *R. mucilaginosa* for pioneer colonizers, *N. sicca* for constitutive species, and *S. exigua* and *L. rimae* for late commensals) and investigated their strain-level single nucleotide variants (SNVs) using *inStrain*. We calculated the non-synonymous to synonymous SNV ratios (N/S ratio) in each community module across the time points (Data S7). The pioneer colonizers and late commensals demonstrated an increasing trend in their N/S ratios (Fig. 7B & 7D) whereas constitutive species showed a slight decreasing trend in N/S ratio (Fig. 7C), suggesting module-specific evolutionary pressures.

We then compared the temporal strain heterogeneity within these species by assessing the population average nucleotide identity (popANI) and shared genome coverage (SGC) at weeks 2 to 4 compared to week 1. The pioneer colonizers (Fig. 7E) and late commensals (Fig. 7G) exhibited greater temporal strain heterogeneity when compared to constitutive species (Fig. 7F), indicating different extents of microevolution towards strain divergence in different modules.

For the non-synonymous SNVs within each species, we also mapped the genes in which they were located to the KEGG Orthology (KO) terms. KO terms with the highest counts of non-synonymous SNVs were visualized and statistically tested to assess whether non-synonymous SNVs were overrepresented in these KO terms when compared to background expectations (Fig. 7H). Except for *P. melaninogenica*, non-synonymous SNVs across all representative species, irrespective of their community module, exhibited significant enrichment in K20386: cylB (ATP-binding cassette, subfamily B, bacterial CylB) and K15558: ophH (phthalate transport system ATP-binding protein). For *P. melaninogenica*, enrichment was observed in K21572: susD, K21573: susC, and K02014: TC.FEV.OM. In addition to K20386: cylB and K15558: ophH, enrichment in K27305: aprG, curF and K24122: ctmAB was observed in *R. mucilaginosa* and *S. exigua*, respectively.

## Discussion

The oral microbiome is involved in various oral diseases and is closely associated with human health. However, investigating the *de novo* assembly of oral microbial communities remains challenging, largely due to the difficulty in defining an accurate starting point for bacterial colonization, given the physiological and anatomical complexity of the oral cavity. Dental implants offer a unique opportunity in this regard. The placement of prosthetic components provides a feasible, well-defined starting point for microbial exposure.

In this study, we applied longitudinal shotgun metagenomic analysis to subgingival plaque samples from peri-implant and adjacent periodontal sites. This approach enabled us to characterize the *de novo* assembly, temporal succession and microevolution of oral microbiome within a complex microenvironment, while also comparing against an adjacent, established community. Our results showed that oral microbiome development is driven by three distinct microbial community modules, each playing a different role in the succession process, with different microevolution trajectories (Fig. 8).

**Fig. 8.**
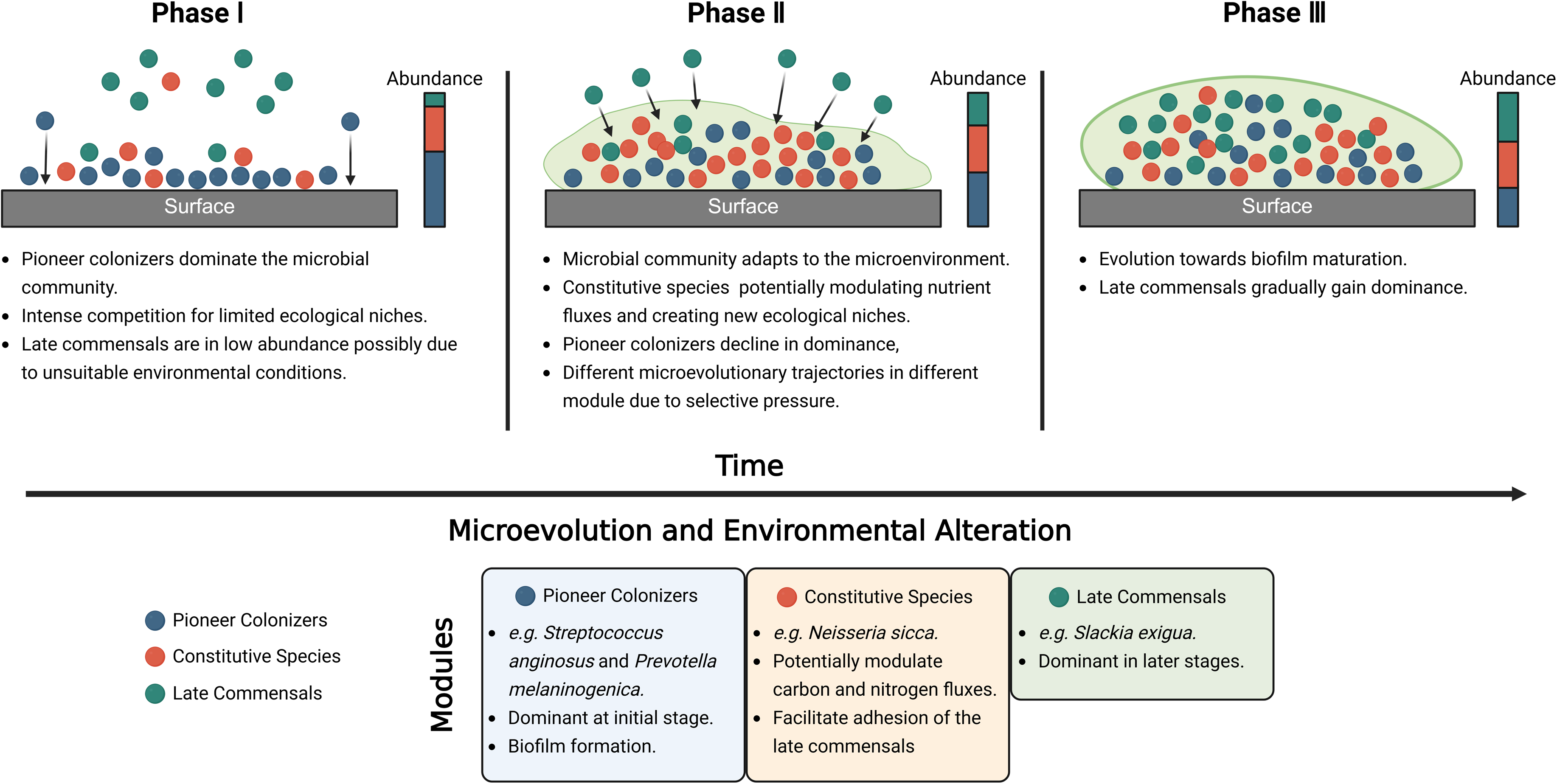
Succession pattern of the oral microbiome in complex microenvironment.

Conventionally, adjacent ecological niches have been considered reservoirs contributing to the formation of the oral microbiome at specific sites [29]. For example, a higher proportion of periodontal pathogens were also detected in peri-implant lesions [30], which suggests a link between peri-implant microbiome and the microbiome from natural teeth. Building on this, our longitudinal cohort revealed distinct taxonomical and functional profiles in peri-implant microbiome compared to the microbiome from adjacent teeth. The two adjacent niches demonstrated significantly different alpha and beta diversity (Fig. 2B & 2C), each harboured a substantial number of exclusive species that were not shared between the two communities (Fig. 2A). These findings suggest that the development of the oral microbiome is not merely a passive consequence of bacterial translocation from adjacent microenvironments but rather involves selective colonization and niche-specific microbial assembly.

The establishment of the microbial community in oral biofilms follows a commonly recognized pattern [31]. It begins with the initial attachment of the pioneer species, followed by the sequential adhesion of secondary and late colonizers [32, 33]. Each stage of colonization modifies the microenvironment, paving the way for the community succession towards maturation of the biofilm. Similarly in our study, distinct abundance trajectories of the microbial species were observed in the peri-implant microbiome (Fig. 5). Based on this, the community was categorized into three modules, namely pioneer colonizers (M1), constitutive species (M2), and late commensals (M3), each comprised of different bacterial species with different functional potentials (Fig. 5).

Previous studies have identified *Streptococcus* species as classical pioneer colonizers in the oral cavity [34–36]. Consistent with these findings, our results also identified multiple *Streptococcus* species in the pioneer module. In addition, we also identified other microbes such as *Prevotella*, *Haemophilus*, and *Rothia* species as potential pioneer colonizers that were essential in the early-stage formation of the peri-implant biofilm. These species were dominant in the initial microenvironment and occupied the ecological niches via exclusive competition. However, they gradually lost their dominance as the microbial community evolved. The high cumulative mutation rates observed in their marker genes may reflect ongoing micro-evolution in concert with the temporal alterations in the microenvironment.

Among the pioneer colonizers, *P. melaninogenica* was a representative being significantly more abundant in peri-implant microbiome compared to periodontal microbiome. *P. melaninogenica* is frequently present in the microbiome of the oral cavity and upper respiratory tract [37], and is potentially associated with periodontitis due to the action of its type □ and □ secretion systems [38, 39]. Interestingly, in our study, we found that *P. melaninogenica* might have different roles to play in periodontal and peri-implant microbiomes, as the same species exhibited significantly higher gene abundance in pathways related to cell wall and extracellular matrix metabolism in peri-implant microbiome than periodontal microbiome, suggesting their importance in biofilm formation (Fig. 4). Similar phenomena were observed in other peri-implant-enriched species including *V. rogosae*, *R. mucilaginosa*, and *S. anginosus*, suggesting that bacterial species may exhibit niche-specific functional adaptations across different ecological sites in the oral cavity.

According to our functional annotations, the pioneer and constitutive species exhibited enrichment in peptidoglycan biosynthesis pathways, the major component of the bacterial cell wall. Peptidoglycan and its fragments generated during the degradation and recycling of the cell wall (e.g. anhydromuropeptides) are multifunctional in the microbial community [27]. Besides maintaining cell wall homeostasis [40], they also act as a community signal to promote biofilm formation [41], and are also responsible for virulence regulation and nutrition complementation in certain bacterial species [41, 42]. Peptidoglycan and its fragments are also major pathogen-associated molecular pattern molecules (PAMPs) and can trigger innate immune responses. These PAMPs can be recognized by TLRs and NLRs thus activating nuclear factor kappa B (NFκB) and mitogen-activated protein kinase (MAPK) intracellular signalling pathways, resulting in the elevation of pro-inflammatory cytokines and promotion of inflammation onset (Fig. 3F). Enrichment in peptidoglycan biosynthesis pathways implies a role for pioneer and constitutive species in not only biofilm formation, but also suggested integration and regulation of the entire microbial community, and more generally, in host-microbiome interactions.

A well-known example of secondary and late colonizers in the oral biofilm is the orange and red complexes discovered by Sorcransky *et al*. [43]. The attachment of orange complex species such as *Fusobacterium nucleatum* increases the complexity of the biofilm and modulates the microenvironment to facilitate the subsequent recruitment of the late colonizers such as *Porphyromonas gingivalis* [44]. This trajectory is typically associated with the transition towards dysbiosis and clinical diseases. Here, we showed that a similar trajectory also exists in the assembly of healthy biofilm, albeit with different microbes (Fig. 5). Represented by *N. sicca*, the constitutive species demonstrated potentials to modulate the microenvironment through pathways like UMP and gondoate biosynthesis (Fig. 5C). UMP biosynthesis is closely linked to cellular carbon–nitrogen metabolic coupling [28] and therefore influences carbon and nitrogen fluxes in the microbial community. The production of gondoate, a monounsaturated fatty acid synthesized independently of oxygen, by constitutive species may enhance membrane fluidity and surface compatibility [45], thereby facilitating the adhesion of late commensals. This was confirmed by our computational simulations based on gLV models (Fig. 6), where the attachment probabilities of late commensal species tended to increase with the presence of constitutive species.

Unlike the other two modules, the late commensals did not demonstrate enrichment in pathways related to cell wall metabolism at week 1 (Fig. 5C). This suggested that they contributed less during the initial biofilm formation. Instead, pathways related to pentapeptide and peptidoglycan biosynthesis began to emerge at week 2 and onwards, possibly marking the proliferation of the late commensals, which is in line with our finding that late commensals were more abundant during weeks 3 and 4 compared to the initial week. Notably the consistently high abundance of coenzyme A biosynthesis across the time points suggested that these species might possess greater metabolic activity compared to other community members, given the central position of coenzyme A in multiple metabolic pathways [46, 47].

Our strain-resolution analyses revealed distinct trajectories of microevolution among the three community modules. Constitutive species demonstrated the most stable evolutionary trajectory, as evidenced by fewer cumulative mutations (Fig. 7A), consistently lower N/S SNVs ratio (Fig. 7C) over time, and robust strain similarity across the time points (Fig. 7F). These findings suggested that constitutive species faced less selective pressure from the peri-implant microenvironment [48] and might play as the “fundamental” components of the peri-implant microbiome. In contrast, pioneer colonizers and late commensals exhibited notable microevolutions with higher cumulative mutation rates (Fig. 7A), increased N/S SNVs ratios over time (Fig. 7B & 7D), and greater temporal strain heterogeneity (Fig. 7E & 7G). These two modules represented divergent responses to the environmental selective process. The pioneer colonizers were dominant during the initial weeks but gradually declined with the succession of peri-implant microbiome, likely due to reduced competitive fitness or niche compatibility, thereby vacating ecological niches. In contrast, late commensals gained dominance over time, potentially facilitated by both environmental modulation and a series of adaptive evolutions. These results are in agreement with previous study showing that in complex microbial communities, initially competitive species can end up in extremely low abundance with the succession process, and that the composition of a stable community was independent of its initial species proportions [49].

The representative species for each community module, except for *P. melaninogenica*, exhibited non-synonymous SNV enrichment in KO terms K20386: cylB (ATP-binding cassette, subfamily B, bacterial CylB) and K15558: ophH (phthalate transport system ATP-binding protein). *CylB* is part of an ATP-binding cassette (ABC) transporter that is responsible for producing and transporting cytolysin, a bacterial virulence factor that disrupts cell membrane and triggers cell lysis [50]. Non-synonymous SNVs in *CylB* can modulate cytolysin secretion, thereby changing the virulence of the microbes. Altered virulence, whether attenuated or enhanced, would reshape host-microbe crosstalk via regulation in immune activation. This indicates how microbes, through genome-level microevolution, dynamically adapted to host-derived pressure, thereby determining which taxa persist and expand as succession proceeds. In parallel, *ophH* is a transporter for phthalates. Under normal conditions, phthalates are unlikely to be abundant in the oral cavity. However, they are used in various dental materials as plasticizers, from resin composites to adhesive cement [51]. After crown cementation, even trace amounts of excess cement can serve as a potential source of leachable phthalates. The enrichment of non-synonymous SNVs in *ophH* further reflected rapid adaptation of the oral microbes to specific chemical compounds present in the oral environment.

Caution should be taken when further interpreting the results of this study, however. Firstly, the oral microbiome is highly individualized [52]. Even the same species can exhibit different phenotypes and functional potentials in different subjects, complicating efforts to assign fixed roles to any single taxon. Secondly, oral ecological niches differ in many aspects such as pH, oxygen tension, nutrient availability, host factors, and surface properties. Consequently, generalizing the succession pattern observed in peri-implant microbiome should be done with care. Future studies investigating evolution at multiple oral sites and explicitly measuring local physicochemical parameters will be important to conclude a universal pattern for oral microbiome development. It should also be noted that the succession of the oral microbiome is a continuous process. However, due to practical constraints, the samples were collected at four discrete time points to infer the community dynamics in this study. Although our findings provide a relatively detailed description of the succession patterns of the peri-implant microbiome, the intervals of sample collection may not fully capture the complexity within this process. Therefore, longer investigations with denser review points are needed for future studies.

## Conclusion

Through this longitudinal cohort study, we showed that oral microbiome development in complex microenvironment is not a passive consequence of bacterial translocation from adjacent niches but a process of selective colonization and niche-specific microbial assembly. Such process follows a structured and dynamic succession pattern driven by three microbial modules, including pioneer colonizers, constitutive species, and late commensals. Each microbial module plays a distinct ecological role and exhibits different trajectories of microevolution. These findings provide a generalizable framework for understanding how oral biofilms originate and evolve in the complex microenvironments of the oral cavity.

## Data availability

Sequences generated during shotgun metagenomics and related metadata were deposited in NCBI Short Reads Archive with Bioproject ID: PRJNA1148759 (https://www.ncbi.nlm.nih.gov/bioproject/?term=PRJNA1148759). Computational codes used for analyses and visualization are available on GitHub (https://github.com/Yuchen-D-Z/Oral_microbiome_development). Other data generated during downstream analyses are provided in the Supplementary Materials.

## Authors’ contributions

Y.Z. designed the study; performed the metagenomic analyses including the taxonomical and functional annotations, the cross-niche and longitudinal comparisons, the bacterial co-occurrence networks, and the genome-level mutation tracking; visualized and interpreted the data; and wrote the manuscript. Y.Y. performed the mathematical work of the gLV model used to predict attachment probability; performed the computational simulation; and contributed to the initial draft. Y.L. collected and stored the clinical samples and curated the raw sequence data as well as subject metadata. E.M.L., D.M., and S.A.N. advised the methodology of the cohort and genome-level mutations, provided the access to computational resources, and revised the manuscript. S.A.N. and Q.Z. supervised the study and raised the fundings. Q.Z. designed the study, conceived and administrated the project. All the authors reviewed, revised, and approved the final manuscript.

## Acknowledgements

We thank the clinicians and nurses at Department of Implant Dentistry, Hospital of Stomatology, Xi’an Jiaotong University for their help in sample collection and storage. We thank King’s Computational Research, Engineering and Technology Environment (CREATE, https://doi.org/10.18742/rnvf-m076) at King’s College London for the computational resources used in metagenomic analyses. We sincerely appreciate the understanding and cooperation of all participants.

## References

1. Mark Welch, J.L., S.T. Ramírez-Puebla, and G.G. Borisy, Oral Microbiome Geography: Micron-Scale Habitat and Niche. Cell Host & Microbe, 2020. 28(2): p. 160–168.

2. Kilian, M., et al., The oral microbiome – an update for oral healthcare professionals. British Dental Journal, 2016. 221(10): p. 657–666.

3. Lamont, R.J., H. Koo, and G. Hajishengallis, The oral microbiota: dynamic communities and host interactions. Nature Reviews Microbiology, 2018. 16(12): p. 745–759.

4. Mark Welch, J.L., et al., Biogeography of a human oral microbiome at the micron scale. Proceedings of the National Academy of Sciences, 2016. 113(6): p. E791–E800.

5. Kolenbrander, P.E., et al., Bacterial interactions and successions during plaque development. Periodontology 2000, 2006. 42(1): p. 47–79.

6. Cugini, C., et al., Dysbiosis From a Microbial and Host Perspective Relative to Oral Health and Disease. Frontiers in Microbiology, 2021. **Volume** 12 **-** 2021.

7. Jorth, P., et al., Metatranscriptomics of the Human Oral Microbiome during Health and Disease. mBio, 2014. 5(2): p. 10.1128/mbio.01012-14.

8. Utter, D.R., C.M. Cavanaugh, and G.G. Borisy, Genome-Centric Dynamics Shape the Diversity of Oral Bacterial Populations. mBio, 2022. 13(6): p. e02414–22.

9. Roodgar, M., et al., Longitudinal linked-read sequencing reveals ecological and evolutionary responses of a human gut microbiome during antibiotic treatment. Genome Res, 2021. 31(8): p. 1433–1446.

10. Cornejo, O.E., et al., Evolutionary and population genomics of the cavity causing bacteria Streptococcus mutans. Mol Biol Evol, 2013. 30(4): p. 881–93.

11. Marsh, P.D. and E. Zaura, Dental biofilm: ecological interactions in health and disease. Journal of Clinical Periodontology, 2017. 44(S18): p. S12–S22.

12. Rosier, B.T., P.D. Marsh, and A. Mira, Resilience of the Oral Microbiota in Health: Mechanisms That Prevent Dysbiosis. Journal of Dental Research, 2018. 97(4): p. 371–380.

13. Hajishengallis, G., R.J. Lamont, and H. Koo, Oral polymicrobial communities: Assembly, function, and impact on diseases. Cell Host & Microbe, 2023. 31(4): p. 528–538.

14. Muguerza-Guevara, K., et al., *In vivo analysis of early biofilm development and cell viability on implant-mimicking abutments at 24 h*, *48 h, and 7 days*. BMC Oral Health, 2025. 25(1): p. 1201.

15. Dieckow, S., et al., *Structure and composition of early biofilms formed on dental implants are complex, diverse, subject-specific and dynamic*. npj Biofilms and Microbiomes, 2024. 10(1): p. 155.

16. Kneaddata - The Huttenhower Lab. Available from: https://huttenhower.sph.harvard.edu/kneaddata/.

17. Truong, D.T., et al., Microbial strain-level population structure and genetic diversity from metagenomes. Genome Res, 2017. 27(4): p. 626–638.

18. Beghini, F., et al., Integrating taxonomic, functional, and strain-level profiling of diverse microbial communities with bioBakery 3. eLife, 2021. 10: p. e65088.

19. Mallick, H., et al., Multivariable association discovery in population-scale meta-omics studies. PLoS Comput Biol, 2021. 17(11): p. e1009442.

20. Grainger, T.N., J.M. Levine, and B. Gilbert, The Invasion Criterion: A Common Currency for Ecological Research. Trends in Ecology & Evolution, 2019. 34(10): p. 925–935.

21. Arnoldi, J.-F., et al., Invasions of ecological communities: Hints of impacts in the invader’s growth rate. Methods in Ecology and Evolution, 2022. 13(1): p. 167–182.

22. Blanco-Miguez, A., et al., Extending and improving metagenomic taxonomic profiling with uncharacterized species with MetaPhlAn 4. bioRxiv, 2022: p. 2022.08.22.504593.

23. Olm, M.R., et al., inStrain profiles population microdiversity from metagenomic data and sensitively detects shared microbial strains. Nature Biotechnology, 2021. 39(6): p. 727–736.

24. Hyatt, D., et al., Prodigal: prokaryotic gene recognition and translation initiation site identification. BMC Bioinformatics, 2010. 11(1): p. 119.

25. Boudreau, M.A., J.F. Fisher, and S. Mobashery, Messenger Functions of the Bacterial Cell Wall-derived Muropeptides. Biochemistry, 2012. 51(14): p. 2974–2990.

26. Müller-Anstett, M.A., et al., Staphylococcal Peptidoglycan Co-Localizes with Nod2 and TLR2 and Activates Innate Immune Response via Both Receptors in Primary Murine Keratinocytes. PLOS ONE, 2010. 5(10): p. e13153.

27. Irazoki, O., S.B. Hernandez, and F. Cava, *Peptidoglycan Muropeptides: Release, Perception,* and Functions as Signaling Molecules. Frontiers in Microbiology, 2019. **Volume** 10 **-** 2019.

28. Braakman, R., et al., Global niche partitioning of purine and pyrimidine cross-feeding among ocean microbes. Science Advances, 2025. 11(1): p. eadp1949.

29. Zhang, Q., et al., Comparison of subgingival and peri-implant microbiome in chronic periodontitis. Chin J Dent Res, 2015. 18(3): p. 155–62.

30. Botero, J.E., et al., Subgingival Microbiota in Peri-Implant Mucosa Lesions and Adjacent Teeth in Partially Edentulous Patients. Journal of Periodontology, 2005. 76(9): p. 1490–1495.

31. Kolenbrander, P.E., et al., Oral multispecies biofilm development and the key role of cell– cell distance. Nature Reviews Microbiology, 2010. 8(7): p. 471–480.

32. Kolenbrander, P.E., et al., Communication among oral bacteria. Microbiology and molecular biology reviews, 2002. 66(3): p. 486–505.

33. Marsh, P., Dental plaque as a microbial biofilm. Caries research, 2004. 38(3): p. 204–211.

34. Sedghi, L., et al., The oral microbiome: Role of key organisms and complex networks in oral health and disease. Periodontology 2000, 2021. 87(1): p. 107–131.

35. Kolenbrander, P.E., Oral microbial communities: biofilms, interactions, and genetic systems. Annual Reviews in Microbiology, 2000. 54(1): p. 413–437.

36. Aruni, A.W., et al., The Biofilm Community-Rebels with a Cause. Curr Oral Health Rep, 2015. 2(1): p. 48–56.

37. Könönen, E. and U.K. Gursoy, Oral Prevotella Species and Their Connection to Events of Clinical Relevance in Gastrointestinal and Respiratory Tracts. Frontiers in Microbiology, 2022. **Volume** 12 **-** 2021.

38. Kondo, Y., et al., Involvement of PorK, a component of the type IX secretion system, in Prevotella melaninogenica pathogenicity. Microbiology and Immunology, 2018. 62(9): p. 554–566.

39. Ibrahim, M., A. Subramanian, and S. Anishetty, Comparative pan genome analysis of oral Prevotella species implicated in periodontitis. Functional & Integrative Genomics, 2017. 17(5): p. 513–536.

40. Salamaga, B., et al., Demonstration of the role of cell wall homeostasis in Staphylococcus aureus growth and the action of bactericidal antibiotics. Proceedings of the National Academy of Sciences, 2021. 118(44): p. e2106022118.

41. Vaidya, S., et al., Bacteria use exogenous peptidoglycan as a danger signal to trigger biofilm formation. Nature Microbiology, 2025. 10(1): p. 144–157.

42. Ruscitto, A., et al., Regulation and Molecular Basis of Environmental Muropeptide Uptake and Utilization in Fastidious Oral Anaerobe Tannerella forsythia. Frontiers in Microbiology, 2017. **Volume** 8 **-** 2017.

43. Socransky, S.S., et al., Microbial complexes in subgingival plaque. J Clin Periodontol, 1998. 25(2): p. 134–44.

44. Kolenbrander, P.E., et al., Bacterial interactions and successions during plaque development. Periodontology 2000, 2006. 42(1).

45. Cronan, J.E., Unsaturated fatty acid synthesis in bacteria: Mechanisms and regulation of canonical and remarkably noncanonical pathways. Biochimie, 2024. 218: p. 137–151.

46. Spry, C., K. Kirk, and K.J. Saliba, Coenzyme A biosynthesis: an antimicrobial drug target. FEMS Microbiol Rev, 2008. 32(1): p. 56–106.

47. Begley, T.P., C. Kinsland, and E. Strauss, The biosynthesis of coenzyme a in bacteria, in Vitamins & Hormones. 2001, Academic Press. p. 157–171.

48. Martiny, J.B.H., et al., Microbial biogeography: putting microorganisms on the map. Nature Reviews Microbiology, 2006. 4(2): p. 102–112.

49. Friedman, J., L.M. Higgins, and J. Gore, Community structure follows simple assembly rules in microbial microcosms. Nature Ecology & Evolution, 2017. 1(5): p. 0109.

50. Shankar, N., et al., Enterococcal cytolysin: activities and association with other virulence traits in a pathogenicity island. International Journal of Medical Microbiology, 2004. 293(7): p. 609–618.

51. Tou, G., et al., Release of leachable products from resinous compounds in the saliva of children with anterior open bite treated with spur. J Appl Oral Sci, 2023. 30: p. e20220227.

52. Belibasakis, G.N., et al., Applications of the oral microbiome in personalized dentistry. Archives of Oral Biology, 2019. 104: p. 7–12.

