## Supplementary Materials for "Development and Genome-level Microevolution of Oral Microbiome during Surface Colonization"

Yuchen Zhang *et al.*

Table S1. Detailed inclusion and exclusion criteria.

| Inclusion Criteria | Exclusion Criteria |
| --- | --- |
| Single tooth implant restoration.  Age between 20 and 60 years.  Well-controlled periodontal conditions.  Adhesive-retained all-ceramic single crown restoration.  A minimum of 4 mm width of keratinized mucosa at the edentulous site. | Presence of progressing periodontitis.  Systemic conditions that might affect bone metabolism or wound healing (e.g. diabetes mellitus or osteoporosis).  Compromised immune conditions.  History of smoking  Antibiotic use within the past six months.  Presence of other dental restorations in the oral cavity.  Pregnancy or lactation. |

**Table S2.** **Demographic features of the cohort.**

| Demographic Features | Number of Participants |
| --- | --- |
| Age in years (mean ± SD) | 37.32 ± 11.22 |
| Gender |  |
| Male (%) | 5 (26.32%) |
| Female (%) | 14 (73.68%) |
| Implant brand |  |
| Bego (%) | 8 (42.11%) |
| Straumann (%) | 6 (31.56%) |
| Hiossen (%) | 3 (15.76%) |
| Zimmer (%) | 1 (5.26%) |
| Osstem (%) | 1 (5.26%) |
| Ethnicity |  |
| Chinese Han (%) | 19 (100%) |

Fig. S1.

Linear models showing significantly higher relative abundance of species *Streptococcus anginosus*, *Prevotella melaninogenica*, *Veillonella rogosae*, *Slackia exigua*, *Neisseria sicca*, *Rothia mucilaginosa*, and *Lancefieldella rimae* in the peri-implant microbiome compared to adjacent periodontal microbiome.

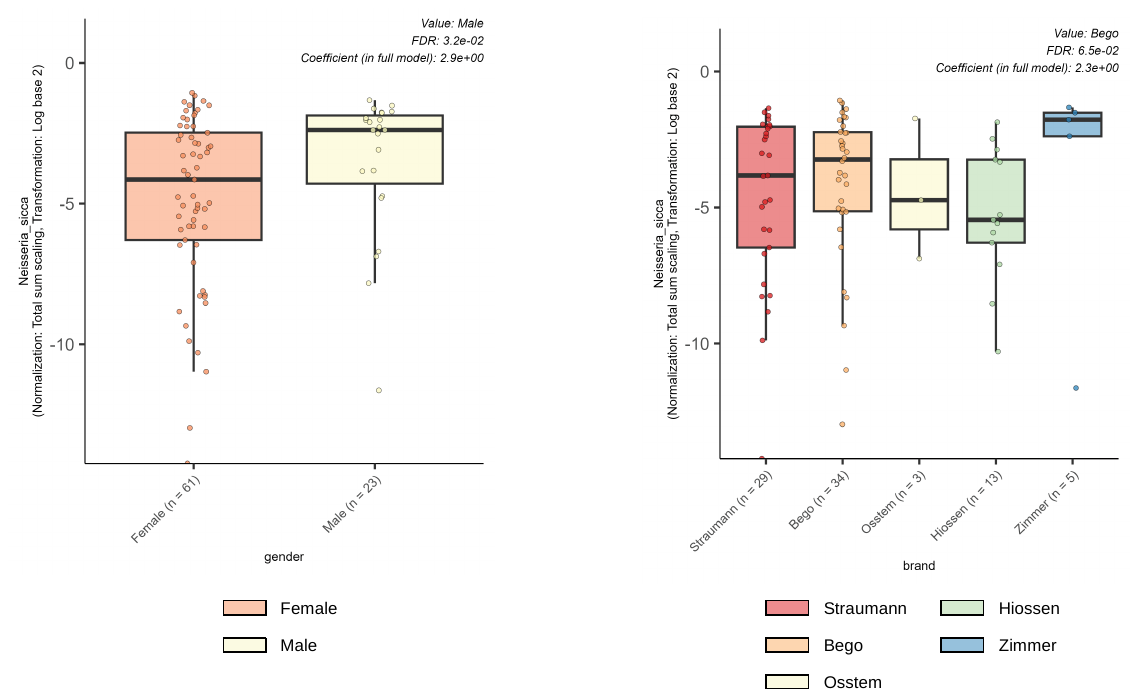

Fig. S2.

Regression models showing that Neisseria *sicca* was positively associated with male gender (p = 0.032) and Bego brand implants (p = 0.065).

Fig. S3.

Linear models showing significantly higher gene abundance related to anhydromuropeptides recycling (PWY-7883 and PWY0-1261) and tRNA processing (PWY0-1479) in the peri-implant microbiome compared to adjacent periodontal microbiome.

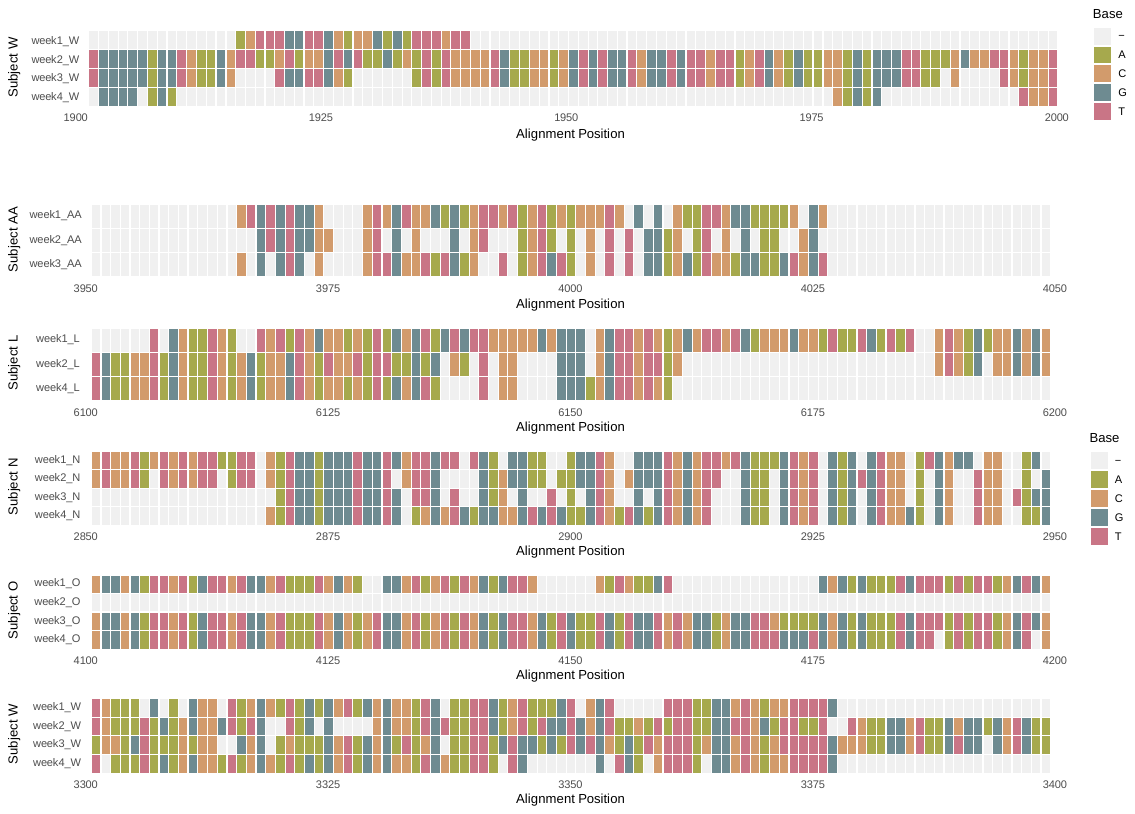

Fig. S4.

*StrainPhlAn* multiple-alignment visualization of mutation hotspots on consensus marker for *Prevotella melaninogenica* (representative species for pioneer colonizers). For each subject, we identified the 100-bp window with the highest single-nucleotide divergence across weeks 1 to 4 using a sliding window (window = 100 bp, step = 50 bp). Tiles show base calls (A, C, G, T) at each alignment position after excluding gaps and ambiguous bases.

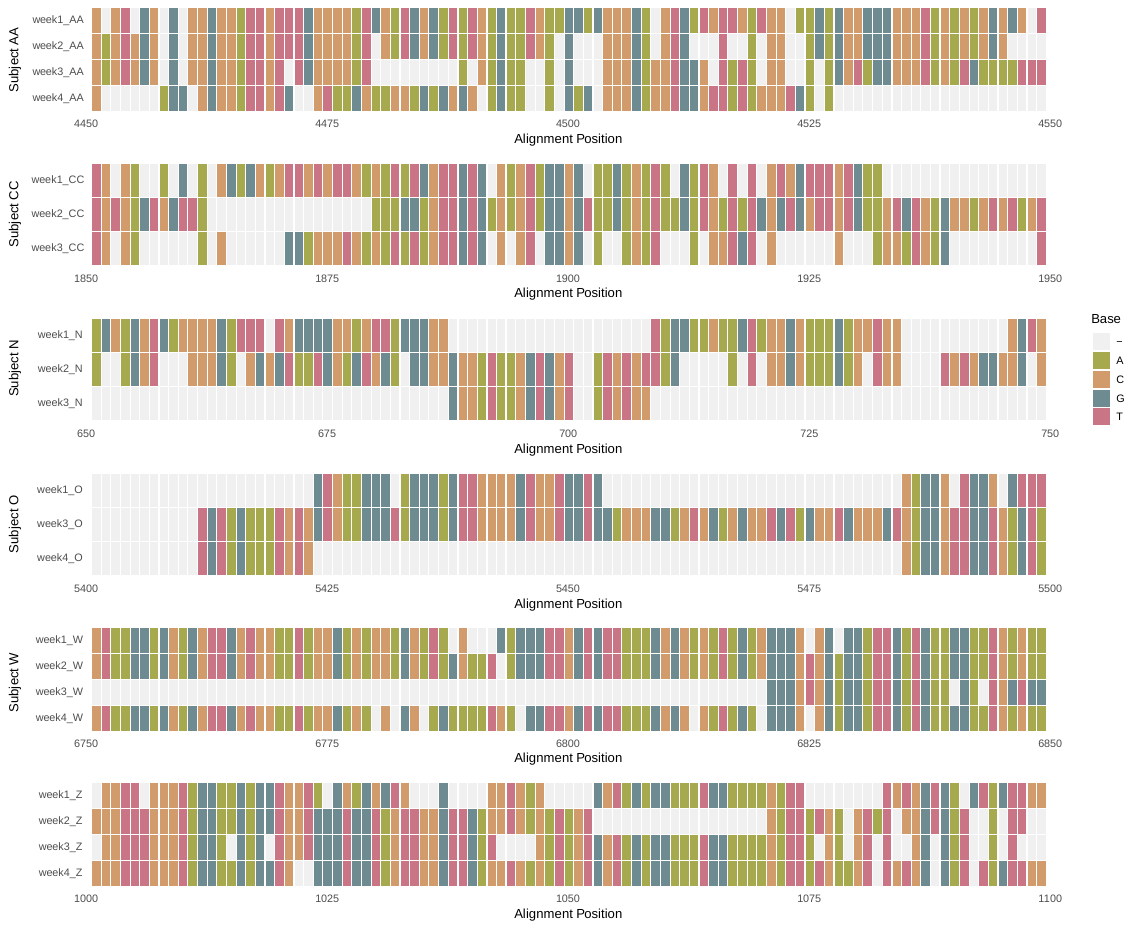

Fig. S5.

Multiple-alignment visualization of mutation hotspots on consensus marker for *Streptococcus anginosus* (representative species for pioneer colonizers).

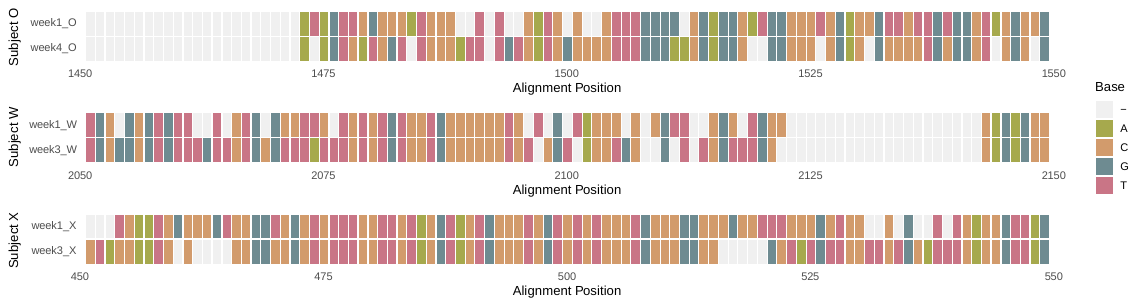

Fig. S6.

Multiple-alignment visualization of mutation hotspots on consensus marker for *Rothia mucilaginosa* (representative species for pioneer colonizers).

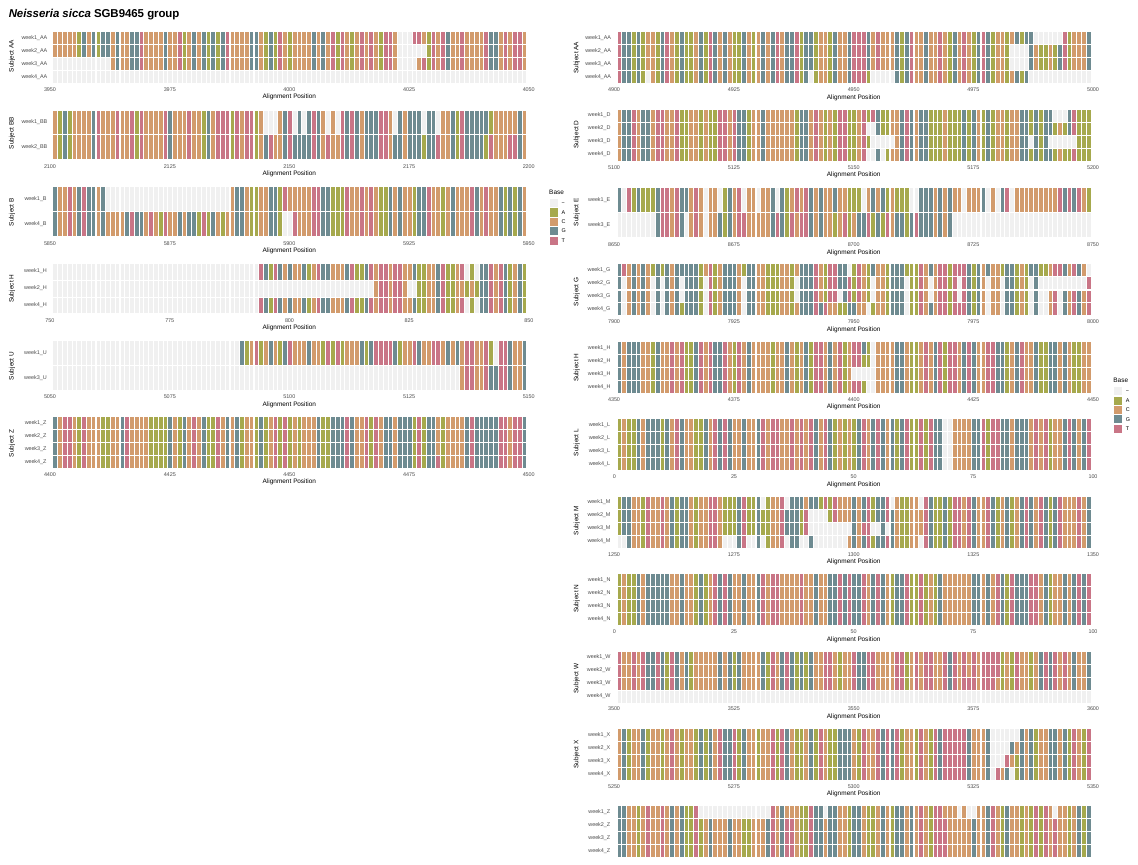

Fig. S7.

Multiple-alignment visualization of mutation hotspots on consensus marker for *Neisseria sicca* (representative species for constitutive species).

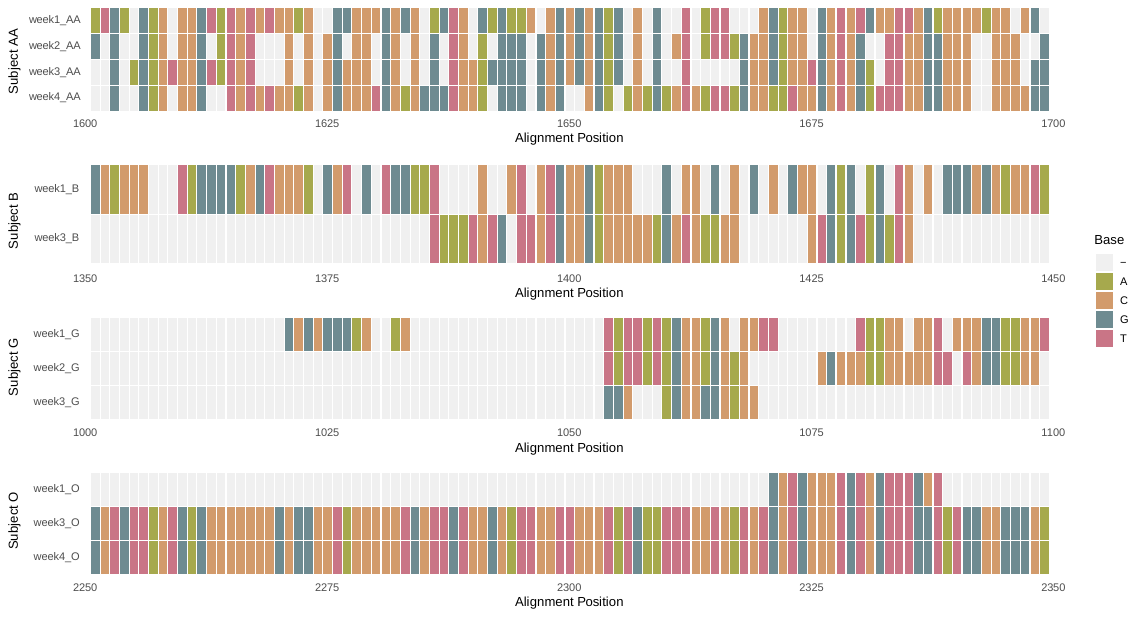

Fig. S8.

Multiple-alignment visualization of mutation hotspots on consensus marker for *Slackia exigua* (representative species for late commensals).

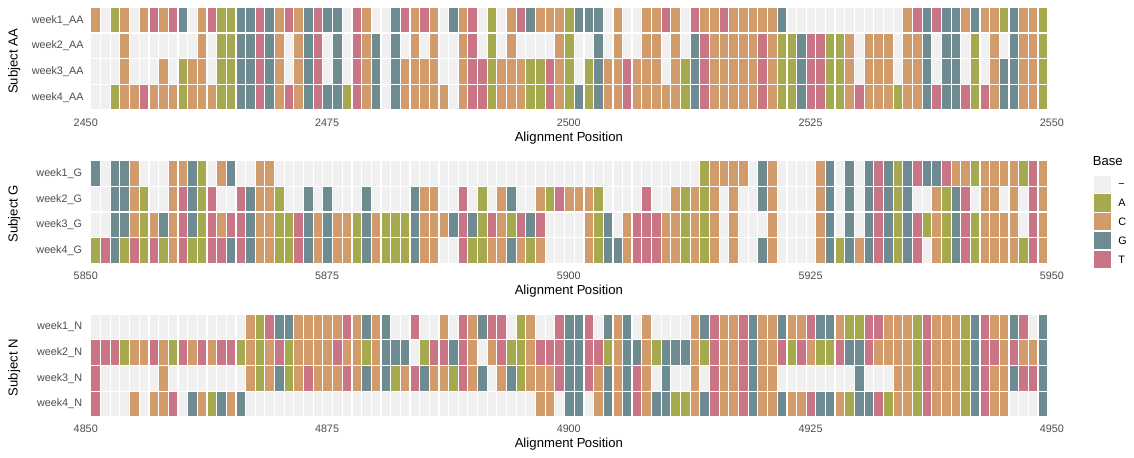

Fig. S9.

Multiple-alignment visualization of mutation hotspots on consensus marker for *Lancefieldella rimae* (representative species for late commensals).

Data S1. (separate file)

The metadata for each sequenced sample.

Data S2. (separate file)

Pathway enrichment or depletion in peri-implant microbiome when compared to the same species in adjacent periodontal microbiome.

Data S3. (separate file)

Members of each community module and their relative abundance.

Data S4. (separate file)

Top 5 enriched pathways in each module at each time point.

Data S5. (separate file)

Spearman correlation coefficients of relative abundances among members within each community module across weeks 1-4 (|rho| > 0.4 and p-FDR < 0.05).

Data S6. (separate file)

Attachment probabilities of late commensal species in the dummy communities P and P + C.

Data S7. (separate file)

Counts of the non-synonymous (N) and synonymous (S) SNVs in each community module across the time points, with N/S ratios.
